# Active Loop Extrusion guides DNA-Protein Condensation

**DOI:** 10.1101/2024.07.03.601883

**Authors:** Ryota Takaki, Yahor Savich, Jan Brugués, Frank Jülicher

**Affiliations:** Max Planck Institute for the Physics of Complex Systems; Max Planck Institute of Molecular Cell Biology and Genetics (MPI-CBG), Dresden, Germany; Center for Systems Biology Dresden, Dresden, Germany; Cluster of Excellence Physics of Life, TU Dresden, Dresden, Germany

## Abstract

The spatial organization of DNA involves DNA loop extrusion and the formation of protein-DNA condensates. While the significance of each process is increasingly recognized, their interplay remains unexplored. Using molecular dynamics simulation and theory we investigate this interplay. Our findings reveal that loop extrusion can enhance the dynamics of condensation and promotes coalescence and ripening of condensates. Further, the DNA loop enables condensate formation under DNA tension and position condensates. The concurrent presence of loop extrusion and condensate formation results in the formation of distinct domains similar to TADs, an outcome not achieved by either process alone.

---

How cells read and process genomic information represents a fundamental question that is not fully understood. This process involves physical interactions between DNA and proteins that transduce sequence information on DNA to express genes and to organize chromatin. Loop extrusion by structural maintenance of chromosomes (SMC) complexes have been identified as a primary candidate of genome organisation and regulation [1–5]. Loop extrusion has been studied through both *in vitro* experiments [6–17] and theoretical approaches [18–25]. DNA loops are involved in the formation of Topologically Associating Domains (TADs) in chromatin. TAD boundaries are determined by the position of CCCTC-binding factor (CTCF) molecules on the DNA [4, 26–31]. Another key process involved in chromatin organization is the formation of biological condensates, a process similar to phase separation [32–40]. Such condensates have been suggested, for example, to play a role in bringing promoters and enhancers into physical proximity [41]. Indeed condensates have been shown to exert capillary forces which could be involved in such processes [42]. These capillary forces are of similar magnitude as forces exerted by SMC molecules during loop extrusion [42]. This raises the question of how loop extrusion and condensate formation synergize to organize chromatin.

In this letter, we use simulation and theory to explore the interplay between loop extrusion and protein condensation in the spacial organization of DNA. We report that DNA loops play a pivotal role in nucleating and positioning protein-DNA co-condensates. The DNA loops not only facilitate the formation of co-condensates but also contribute to their stability under mechanical tension along DNA. We further discuss how loop extrusion and condensation contribute to the emergence of domains in chromatin contact maps, which characterize the DNA spatial organization.

Here we consider a configuration often used in biophysical studies of DNA, where a single DNA molecule is attached at both ends to a surface [7, 42] (Fig.1). We perform Langevin dynamics simulations that incorporate three distinct types of particle representing DNA segments, proteins, and SMC molecules. These particles interact according to Lennard-Jones potentials as well as FENE potentials along the DNA contour [43]. DNA-Protein and Protein-Protein interactions are attractive while the interaction among DNA segments is repulsive. DNA segments are coarse-grained by particles with 10 nm diameter, and all three particle types have the same diameter. SMC molecules can bind to the DNA strand in the region indicated in blue in Fig.1. Upon binding of an SMC molecule to a DNA segment, a two-sided loop extrusion process is initiated. The loop extrusion process terminates when the extruded DNA reaches a boundary of the blue region, mimicking the role of CTCF molecules [26, 44]. We start our simulation from randomized initial positions of proteins and DNA segments, with DNA ends fixed at a prescribed distance *L*_end_ *µ*m. The contour length of DNA is set to *L*_*c*_ = 16.5 *µ*m, corresponding to λ-DNA [42]. See Supplementary Information (SI) for simulation details.

**FIG 1.**
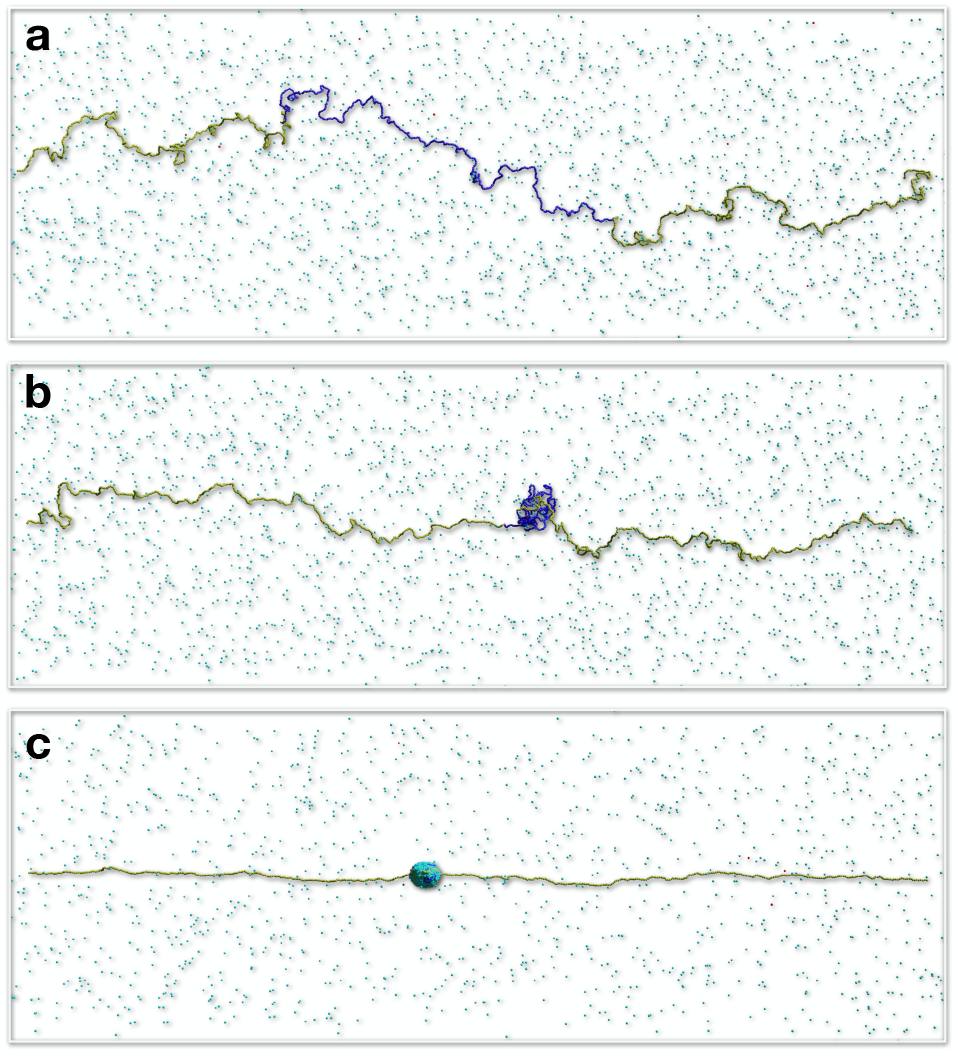
(a) Initial configuration of the simulation with yellow and blue DNA particles indicating the DNA strand and SMC binding sites on DNA, respectively. Light blue particles represent proteins, and red particles SMC molecules. (b) Example of a DNA loop created by an SMC molecule. (c) Example of a Protein-DNA co-condensate. (a)-(c) are independent simulations for *L*_end_ = 6 *µ*m.

We first focus on the dynamics of condensate growth. Similar to conventional droplet kinetics, DNA-protein condensates grow through coalescence and Ostwald ripening when multiple condensates exist. Fig.2a displays the size *S* of the largest droplet as a function of simulation time *t*, averaged over three simulation trajectories. The size *S* is defined as the number of DNA and protein particles contained in the condensate (section IV.A in the SI). Step-wise increases of *S*(*t*) indicate coalescence events, while gradual growth corresponds to ripening. We compare simulations without loop extrusion (Fig.2a, left) to simulations with loop extrusion (Fig.2a, right) for different *L*_end_. This comparison reveals that loop extrusion accelerates ripening but also enhances coalescence.

**FIG 2.**
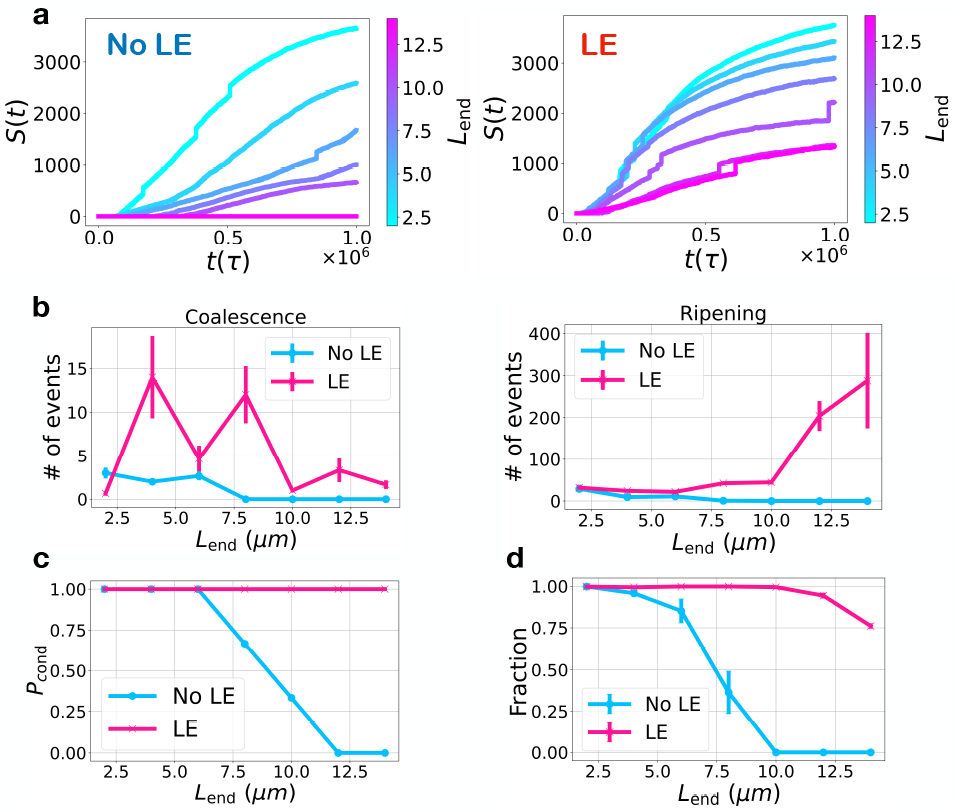
(a) Size of the largest condensates *S*(*t*) as a function of time *t*. Here *τ* is the unit-time in our simulation (SI). The color bars represent the DNA end-to-end distance, *L*_end_. Results without loop extrusion (No LE) and with loop extrusion (LE) are shown. (b) Number of condensate disappearance events due to droplet coalescence (left) or disassembly during ripening (right). Lines represent scenarios without loop extrusion (blue) and with loop extrusion (red). (c) Probability of condensate presence in the final frame of the simulation. (d) Fraction of SMC-binding DNA segments (blue in Fig.1) within the condensate. Error bars indicate the standard error across three simulations with same parameter values.

To provide further insight into the condensate growth dynamics, we count the number of events where condensates disappear either by coalescence or by Ostwald ripening, see Fig.2b. We find a systematic increase of condensate coalescence events in the presence of loop extrusion as compared to the absence of loop extrusion. Furthermore we observe that droplet disappearance by ripening is strongly enhanced for large *L*_end_. The enhancement of Ostwalt ripening by loop extrusion can be understood as follows: condensates outside the loop are subjected to the tension of the polymer and therefore disfavored as compared to the condensate inside the loop which is not subject to tension (section II in the SI).

We next calculated the probability of condensate formation (*P*_cond_), defined as the probability of condensate formation in the final frame of our simulation trajectories (Fig. 2c). For short *L*_end_, a condensate is present irrespective of whether a loop was extruded. For larger *L*_end_ the probability to find a condensate drops and eventually vanishes if no loop is extruded, similar to previous results [42]. Interestingly, loop extrusion enables condensate formation even at large *L*_end_. Loop extrusion can also position condensates. In fact, in our simulations, condensates are often found where the loop is formed. To quantify this co-localization, we compute in Fig.2d the fraction of SMC-binding DNA segments (blue in Fig.1) within the condensate. With loop extrusion, these segments are largely included inside the condensate, even for larger values of *L*_end_. In contrast, without loop extrusion, this fraction decreases with increasing *L*_end_.

The positioning of condensates by loop extrusion can be discussed by considering three possible scenarios. In scenario (a), the DNA loop is located outside of the condensate (Fig.3a). In the scenario (b), the condensate is located at the DNA loop and only the DNA loop is inside the condensate (Fig.3b). Finally in scenario (c) the DNA loop is inside the condensate together with additional DNA segments of length *δ* (Fig.3c). Our simulations suggest that condensation within the DNA loop is not affected by mechanical tension. Therefore condensates form reliably in the scenario (b) and (c) containing the loop. To understand the effect of DNA loop on the formation of condensates, we use a simple model of co-condensation [42]. In this model, the free energy of the configuration is the sum, *F* = *F*_*d*_ + *F*_*p*_, where

**FIG 3.**
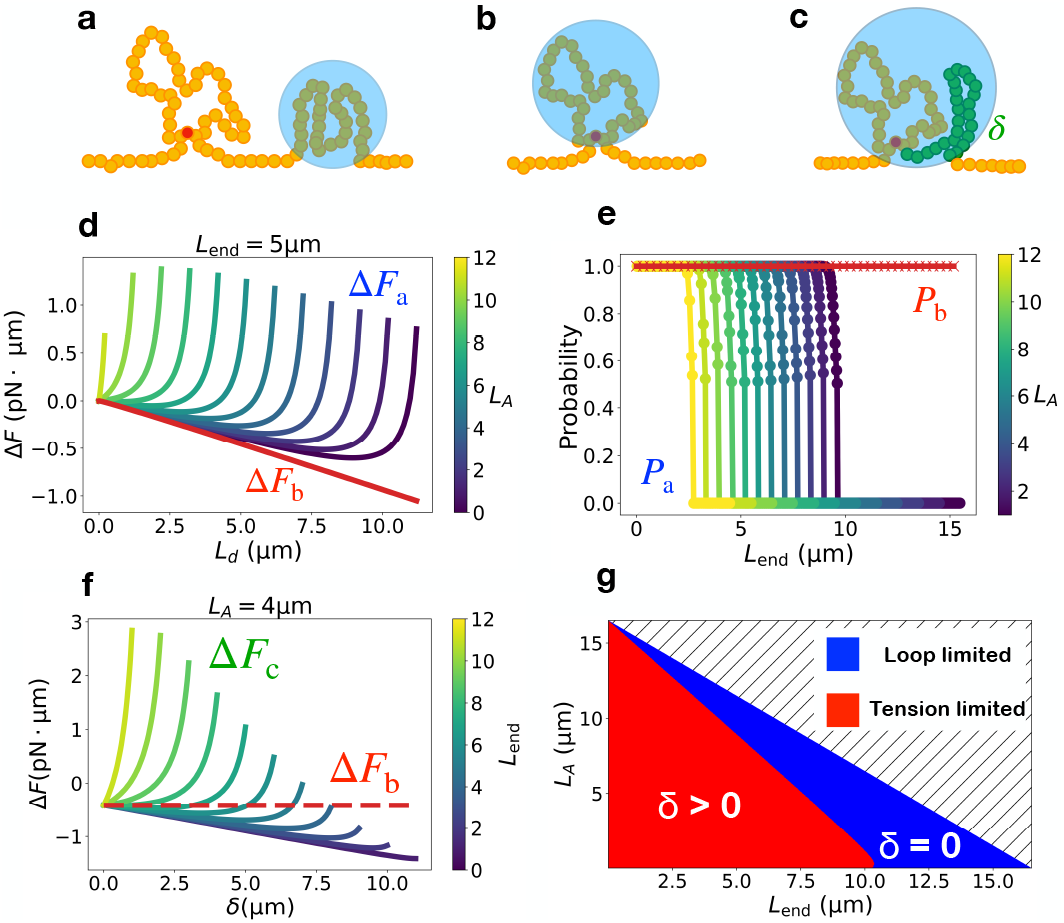
(a)-(c) Schematic representations of three scenarios of protein-DNA co-condensation. Yellow, blue, and red circle indicates DNA segments, condensate, and SMC protein, respectively. (a) Condensate positioned outside the DNA loop. (b) Condensates at the DNA loop. (c) Condensate at the loop, including extra DNA length *δ* (green). (d) Free energy, Δ*F*_*a*_, for scenario (a) compared to Δ*F*_*b*_ corresponding to (b), as a function of the length *L*_*d*_ within the condensate for different loop length *L*_*A*_. (e) Probability *P*_*a*_ (*P*_*b*_) of condensation formation for scenario (a) (scenario (b)) as a function of *L*_end_. (f) Free energy Δ*F*_*c*_ for scenario (c) as a function of *δ* compared to Δ*F*_*b*_ for different *L*_end_. (g) Phase diagram of condensation in the presence of DNA loop as a function of *L*_*A*_ and *L*_end_. The loop limited regime (blue) and the tension limited regime (red) are indicated. The hatched region is physically inaccessible.

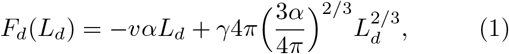

is the free energy of the condensate and

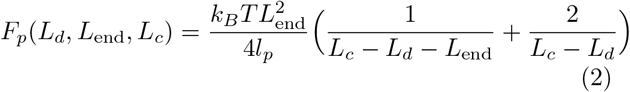

is the free energy of the non-condensed DNA. Here *L*_*d*_ is the length of the DNA segments inside the condensate, *k*_*B*_ is the Boltzmann constant, *T* represents the temperature, and *l*_*p*_ is the persisten ce length of DNA. The parameters *α, γ*, and *v* are the inverse of the DNA packing density, surface tension of the condensate, and condensation free energy per volume, respectively [42]. Below we use parameters values obtained for forkhead box protein A1 [42]: *α* = 0.04 *µ*m^2^, *γ* = 0.04 pN *µ*m^−1^, *v* = 2.6 pN *µ*m^−2^ and *l*_*p*_ = 50 nm.

We first consider scenario (a) (Fig.3a). In this case the condensate is located outside the DNA loop of length LA. We define the energy difference Δ*F*_*a*_ = *F*_*d*_(*L*_*d*_) + *F*_*p*_(*L*_*d*_, *L*_end_, *L*_*c*_ ™ *L*_*A*_) ™ *F*_*p*_ (0, *L*_end_, *L*_*c*_ ™ *L*_*A*_) as the difference of the free energy before and after condensate formation, where we have taken into account that the DNA contour length *L*_*c*_ is effectively reduced by the loop length *L*_*A*_. The stable condensate size is then obtained by minimizing Δ *F*_*a*_ with respect to *L*_*d*_.

In scenario (b), the condensate c length *L*_*d*_ = *L*_*A*_ and the free energy cange due to con-densate formation is simply given by Δ*F*_*b*_ = *F*_*d*_(*L*_*A*_) Correspondingly, in scenario (c), we define Δ*F*_*c*_ = *F*_*d*_(*L*_*A*_ + *δ*) + *F*_*p*_(*L*_*A*_ + *δ, L*_end_, *L*_c_) − *F*_*p*_(*L*_*A*_, *L*_end_, *L*_c_). When *δ* = 0, the DNA length inside the condensate equals to the loop length *L*_*A*_, reducing this case to scenario (b).

We show Δ*F*_*a*_ in Fig.3d, as well as Δ*F*_*b*_ for *L*_end_ = 5 *µ*m. Increasing the loop length *L*_*A*_ shifts the minimum position of Δ*F*_*a*_ towards smaller values of *L*_*d*_ until the condensate vanishes via a first-order phase transition, similar to the one reported previously [42]. Fig.3d reveals that Δ*F*_*b*_ < Δ*F*_*a*_, implying that a condensate will always form inside the loop, irrespective of the tension on the DNA. Thus, the DNA loop guides condensate formation. Fig.3e, shows the conditional probability of generating a condensate outside the loop (*P*_*a*_) as a function of *L*_end_. Increasing *L*_*A*_ shifts the curves for *P*_*a*_ towards smaller *L*_end_ values: the tension induced by the DNA loop narrows the range where condensation outside the loop is possible. However condensates in the loop are always favored (*P*_*b*_ = 1).

Fig.3f shows the free energy profile Δ*F*_*c*_ as a function of *δ* together with Δ*F*_*b*_. This shows that for sufficiently large *L*_end_ the free energy is minimal for *δ* = 0 and the condensate size is equal to the loop size. We call this loop limited condensate. As *L*_end_ is decreased below the critical value 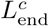, *F*_*c*_ exhibits a minimum at *δ >* 0 corresponding to a condensate containing a DNA segment that is longer than the loop. In this regime the condensate is tension limited. At 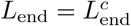 where *δ* vanishes, a continuous second order transition occurs. Fig.3g shows the phase diagram for condensates in the presence of a loop. In the blue region, *δ* = 0 and condensates are loop limited. In the red region, *δ >* 0 and condensates are tension limited. Both regions meet at a second order transition line.

We now discuss the effects of loops on DNA conformations inside the condensates. Both condensation and loop extrusion lead to a local accumulation of DNA segments, see Fig. 1. Such accumulation fosters contacts between DNA segments, even when they are at a distance along the sequence. The probabilities of such contacts are characterized by a contact map, a key tool to understand chromatin organization [45–49]. We determine contact maps for different end-to-end distance in our simulations. Fig.4 presents examples of contact maps with contact probability between two segments *i* and *j*, where *i* and *j* are the DNA particle indices of two DNA segments. In Fig. 4, contact maps are shown for DNAprotein co-condensation without loop extrusion (left column), for loop extrusion without condensation (middle column), and for both condensation and loop extrusion (right column), for different values of *L*_end_. We find that if only condensation happens, contact maps show many but irregularly positioned disordered contacts for short *L*_end_. These contacts disappear as *L*_end_ is increased and condensates dissolve. For only loop extrusion, contacts occur within the loop and are dominated by short-ranged contacts. When condensation and loop extrusion are combined, square patterns of contacts resembling TADs emerge. These square patterns imply that contacts over longer distances along the chain are prominent.

**FIG 4.**
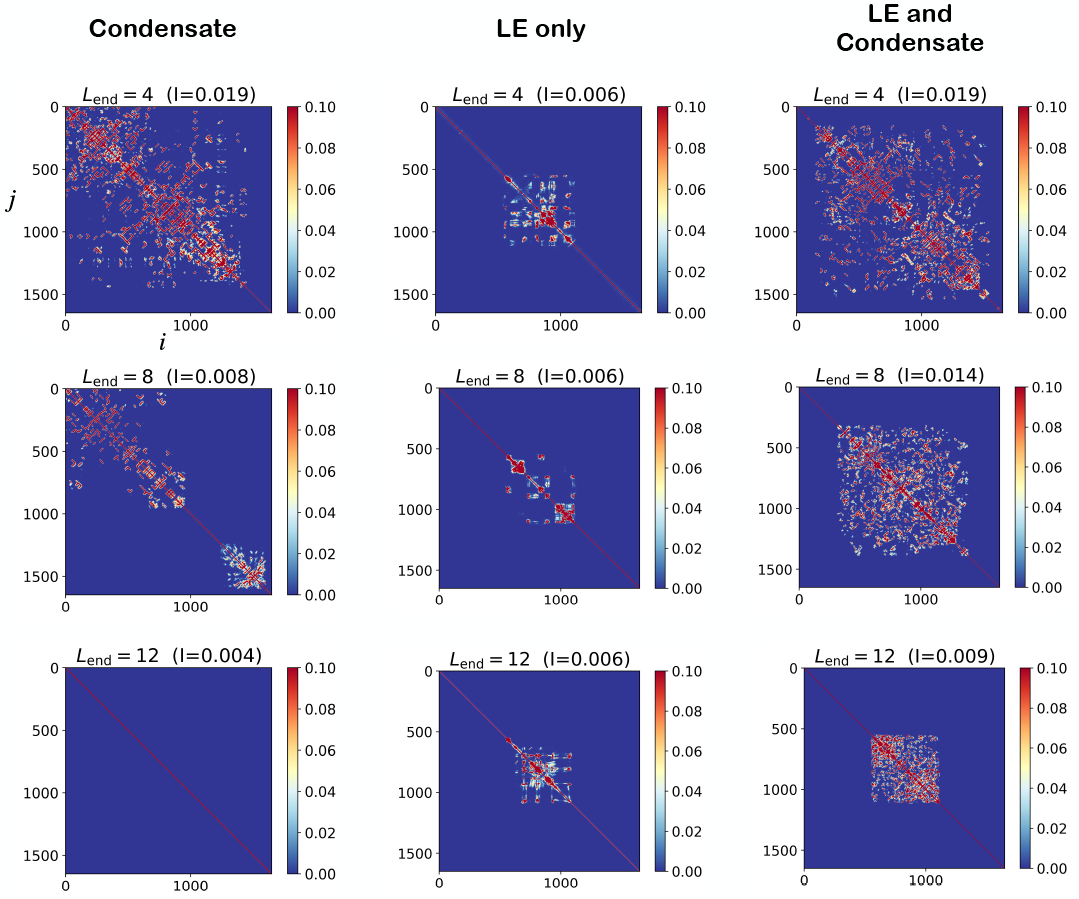
Contact maps obtained in simulations for only condensation (left), only loop extrusion (middle), and both loop extrusion and condensation (right) as a function of DNA particle indices *i* and *j*. Contact maps are presented for *L*_end_ = 4, 8, 12 *µ*m (top to bottom). The contact probability between particle *i* and *j* is shown as a color code. The fraction *I* of contacts, defined as the sum of the all contact probabilities normalized by the total number of pairs (1650^2^), is given.

To further characterise the structures generated by condensation and loop extrusion, we consider the polymeric configuration of loops and condensates. Fig.5a shows example configurations obtained in simulations: a DNA loop without condensation (left) and a DNA loop within a condensate (right). This reveals a coil-like structure of the DNA loop and a densely packed DNA within the condensate. Fig.5b shows, the radius of gyration (*R*_*g*_) of the DNA loop as a function of loop length, exhibiting a scaling behavior with an exponent *ν* = 0.54, similar to the Flory exponent *ν* = 3*/*5 of a polymer in a good solvent [43]. In contrast, *R*_*g*_ of the DNA in exhibits different scaling behavior with is close to the exponent *ν* = 1*/*3 of a c in a poor solvent [43].

**FIG 5.**
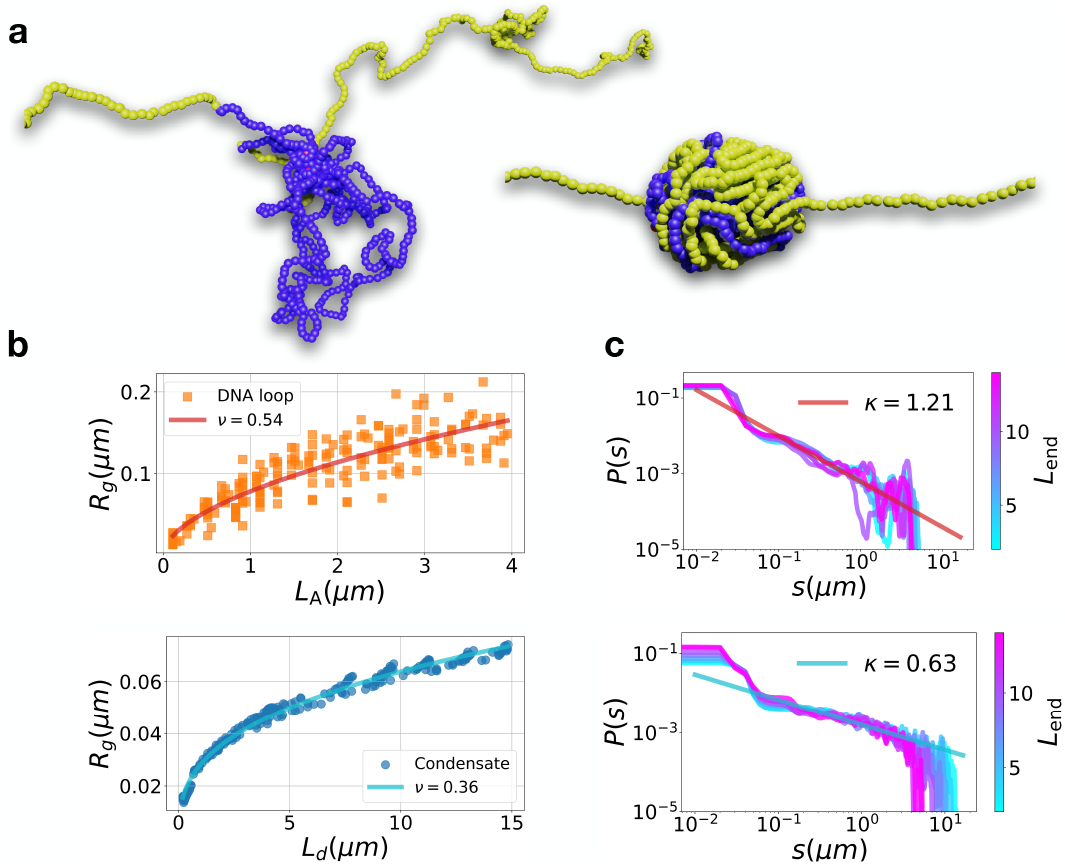
(a) Representative snapshots showing a DNA loop without condensation (left) and a DNA-Protein condensate containing the DNA loop (right). Yellow and blue particles indicate the DNA strand and SMC binding site on DNA, respectively. Protein particles are not shown to emphasize the DNA structure. *L*_end_ = 6 *µm*. (b) Radius of gyration *R*_*g*_ of DNA loops without condensation as a function loop length *L*_*A*_ (top, *ν* ≃ 0.54) and of DNA within a condensate as a function of condensed DNA length *L*_*d*_ (bottom, *ν* ≃ 0.36). The data is fit using *R*_*g*_ ∼ *L*^*ν*^ (solid lines). (c) Probability of contact, *P*, as a function of the DNA length between contacts, *s*, for different *L*_end_ for loop extrusion without condensation (top, *κ* = 1.21) and for loop extrusion with condensation (bottom, *κ* = 0.63). The data is fit using *P* ∼ *s*^−*κ*^ (solid lines). The reported values of *κ* are the mean value of the exponents for different *L*_end_ fit in the range 5 · 10^−2^ *µm* < *s* < 3 *µm*.

Using these results, we can explain the difference between the contact maps shown in Fig.4. In condensation, the looped polymer behaves as a random coil favoring short range contacts. In probability decreases for increasing di polymer, giving rise to the short rang shown in Fig.4, middle. For a loop in sate, the contact probability remains hi the longer ranged contact map. We us ment for the scaling of the contact pro as a function of the DNA length *s* betw points. Assuming a probability of cont Using these results, we can explain the difference be-tween the contact maps shown in Fig.4. In the absence of condensation, the looped polymer behaves as a random coil favoring short range contacts. Indeed, the contact probability decreases for increasing distance along the polymer, giving rise to the short ranged contact maps shown in Fig.4, middle. For a loop inside the condensate, the contact probability remains high, giving rise to the longer ranged contact map. We use a simple argument for the scaling of the contact probability P ∼ s − κ as a function of the DNA length s between the contacts points. Assuming a probability of contacts proportional to the squared density, we have

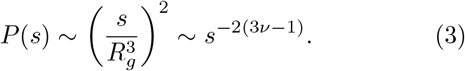

The Flory exponent of *ν* = 3*/*5 for a good solvent implies *κ* = 8*/*5 while for a poor solvent *ν* = 1*/*3 leads to *κ* = 0.

We tested this scaling argument in our simulation. Fig.5c (top) shows *P* (*s*) for a loop without condensate. We find *κ* ≃ 1.21, which is in good agreement with the estimate *κ* = 1.24 using *κ* = 6*ν* − 2 with *ν* = 0.54 (Fig.5b, top). For loop extrusion with condensation, we obtained *κ* ≃ 0.63 (Fig.5c, bottom). This is longer ranged as compared to the case without condensate, but deviates from the estimate *κ* = 0.16 using *ν* = 0.36 (Fig.5b, bottom). This difference may stem from finite-size and finite-time effects in our simulations. However, the change of the exponents for loops with (*κ* = 0.63) and without condensation (*κ* = 1.21) is significant, and highlights the longer range contacts in a condensed DNA loop.

In summary, we investigated the interplay between DNA loop extrusion and DNA-protein co-condensation. We found that loop extrusion stabilises DNA-protein co-condensation under tension and positions the condensate on DNA. We identified a regime of loop limited condensates, where condensates size is set by the DNA loop size. Our work shows that by combining loop extrusion and condensation, a DNA organization with the characteristics of TADs naturally emerges. In the absence of the condensation, the resulting contact maps remain short ranged. The condensation facilitates the close contacts of distant DNA segments within the TAD. Our work underscores chromatin organization in the nucleus may emerge from the interplay between loop extrusion and protein-DNA co-condensation.

## References

[1] K. Nasmyth and C. H. Haering, Cohesin: its roles and mechanisms, Annual review of genetics 43, 525 (2009).

[2] E. Kim, R. Barth, and C. Dekker, Looping the genome with smc complexes, Annual Review of Biochemistry 92 (2023).

[3] I. F. Davidson and J.-M. Peters, Genome folding through loop extrusion by smc complexes, Nature reviews Molecular cell biology 22, 445 (2021).

[4] S. Yatskevich, J. Rhodes, and K. Nasmyth, Organization of chromosomal dna by smc complexes, Annual review of genetics 53, 445 (2019).

[5] C. Hoencamp and B. D. Rowland, Genome control by smc complexes, Nature Reviews Molecular Cell Biology, 1 (2023).

[6] T. Terakawa, S. Bisht, J. M. Eeftens, C. Dekker, C. H. Haering, and E. C. Greene, The condensin complex is a mechanochemical motor that translocates along dna, Science 358, 672 (2017).

[7] M. Ganji, I. A. Shaltiel, S. Bisht, E. Kim, A. Kalichava, C. H. Haering, and C. Dekker, Real-time imaging of dna loop extrusion by condensin, Science 360, 102 (2018).

[8] S. Golfier, T. Quail, H. Kimura, and J. Brugués, Cohesin and condensin extrude dna loops in a cell cycle-dependent manner, Elife 9, e53885 (2020).

[9] E. Kim, A. M. Gonzalez, B. Pradhan, J. van der Torre, and C. Dekker, Condensin-driven loop extrusion on supercoiled dna, Nature Structural & Molecular Biology 29, 719 (2022).

[10] E. Kim, J. Kerssemakers, I. Shaltiel, C. Haering, and C. Dekker, Dna-loop extruding condensin complexes can traverse one another, Biophysical Journal 118, 380a (2020).

[11] B. Pradhan, T. Kanno, M. Umeda Igarashi, M. S. Loke, M. D. Baaske, J. S. K. Wong, K. Jeppsson, C. Björkegren, and E. Kim, The smc5/6 complex is a dna loop-extruding motor, Nature, 1 (2023).

[12] I. F. Davidson, B. Bauer, D. Goetz, W. Tang, G. Wutz, and J.-M. Peters, Dna loop extrusion by human cohesin, Science 366, 1338 (2019).

[13] Y. Kim, Z. Shi, H. Zhang, I. J. Finkelstein, and H. Yu, Human cohesin compacts dna by loop extrusion, Science 366, 1345 (2019).

[14] J.-K. Ryu, S.-H. Rah, R. Janissen, J. W. Kerssemakers, A. Bonato, D. Michieletto, and C. Dekker, Condensin extrudes dna loops in steps up to hundreds of base pairs that are generated by atp binding events, Nucleic acids research 50, 820 (2022).

[15] B. Pradhan, R. Barth, E. Kim, I. F. Davidson, B. Bauer, T. van Laar, W. Yang, J.-K. Ryu, J. van der Torre, J.-M. Peters, et al., Smc complexes can traverse physical roadblocks bigger than their ring size, Cell reports 41 (2022).

[16] J.-K. Ryu, A. J. Katan, E. O. van der Sluis, T. Wisse, R. de Groot, C. H. Haering, and C. Dekker, The condensin holocomplex cycles dynamically between open and collapsed states, Nature structural & molecular biology 27, 1134 (2020).

[17] G. Shi, L. Liu, C. Hyeon, and D. Thirumalai, Interphase human chromosome exhibits out of equilibrium glassy dynamics, Nature communications 9, 3161 (2018).

[18] R. Takaki, A. Dey, G. Shi, and D. Thirumalai, Theory and simulations of condensin mediated loop extrusion in dna, Nature Communications 12, 5865 (2021).

[19] E. J. Banigan and L. A. Mirny, Limits of chromosome compaction by loop-extruding motors, Physical Review X 9, 031007 (2019).

[20] E. J. Banigan and L. A. Mirny, Loop extrusion: theory meets single-molecule experiments, Current opinion in cell biology 64, 124 (2020).

[21] E. J. Banigan and L. A. Mirny, The interplay between asymmetric and symmetric dna loop extrusion, Elife 9, e63528 (2020).

[22] B. Chan and M. Rubinstein, Theory of chromatin organization maintained by active loop extrusion, Proceedings of the National Academy of Sciences 120, e2222078120 (2023).

[23] J. F. Marko, P. De Los Rios, A. Barducci, and S. Gruber, Dna-segment-capture model for loop extrusion by structural maintenance of chromosome (smc) protein complexes, Nucleic acids research 47, 6956 (2019).

[24] A. Bonato and D. Michieletto, Three-dimensional loop extrusion, Biophysical Journal 120, 5544 (2021).

[25] B. Chan and M. Rubinstein, Activity-driven chromatin organization during interphase: compaction, segregation, and entanglement suppression, Proceedings of the National Academy of Sciences 121, e2401494121 (2024).

[26] I. F. Davidson, R. Barth, M. Zaczek, J. van der Torre, W. Tang, K. Nagasaka, R. Janissen, J. Kerssemakers, G. Wutz, C. Dekker, et al., Ctcf is a dna-tension-dependent barrier to cohesin-mediated loop extrusion, Nature, 1 (2023).

[27] D. Jeong, G. Shi, X. Li, and D. Thirumalai, Structural basis for the preservation of a subset of topologically associating domains in interphase chromosomes upon cohesin depletion, Elife 12, RP88564 (2024).

[28] W. Schwarzer, N. Abdennur, A. Goloborodko Pekowska, G. Fudenberg, Y. Loe-Mie, N. A. Fonseca, W. Huber, C. H. Haering, L. Mirny, et al., Two independent modes of chromatin organization revealed by cohesin removal, Nature 551, 51 (2017).

[29] G. Fudenberg, M. Imakaev, C. Lu, A. Goloborodko, N. Abdennur, and L. A. Mirny, Formation of chromo-somal domains by loop extrusion, Cell reports 15, 2038 (2016).

[30] E. J. Banigan, W. Tang, A. A. van den Berg, R. R. Stocsits, G. Wutz, H. B. Brandao, G. A. Busslinger, J.-M. Peters, and L. A. Mirny, Transcription shapes 3d chromatin organization by interacting with loop extrusion, Proceedings of the National Academy of Sciences 120, e2210480120 (2023).

[31] A. Dey, G. Shi, R. Takaki, and D. Thirumalai, Structural changes in chromosomes driven by multiple condensin motors during mitosis, Cell Reports 42 (2023).

[32] A. A. Hyman, C. A. Weber, and F. Jülicher, Liquid-liquid phase separation in biology, Annual review of cell and developmental biology 30, 39 (2014).

[33] K. Rippe, Liquid–liquid phase separation in chromatin, Cold Spring Harbor perspectives in biology 14, a040683 (2022).

[34] J. T. King and A. Shakya, Phase separation of dna: From past to present, Biophysical Journal 120, 1139 (2021).

[35] K. Shrinivas, B. R. Sabari, E. L. Coffey, I. A. Klein, A. Boija, A. V. Zamudio, J. Schuijers, N. M. Hannett, P. A. Sharp, R. A. Young, and A. K. Chakraborty, Enhancer features that drive formation of transcriptional condensates, Molecular Cell 75, 549 (2019).

[36] M.-T. Wei, Y.-C. Chang, S. F. Shimobayashi, Y. Shin, A. R. Strom, and C. P. Brangwynne, Nucleated transcriptional condensates amplify gene expression, Nature cell biology 22, 1187 (2020).

[37] J. E. Henninger, O. Oksuz, K. Shrinivas, I. Sagi, G. LeRoy, M. M. Zheng, J. O. Andrews, A. V. Zamudio, C. Lazaris, N. M. Hannett, et al., Rna-mediated feed-back control of transcriptional condensates, Cell 184, 207 (2021).

[38] J. A. Morin, S. Wittmann, S. Choubey, A. Klosin, S. Golfier, A. A. Hyman, F. Jülicher, and S. W. Grill, Sequence-dependent surface condensation of a pioneer transcription factor on dna, Nature Physics 18, 271 (2022).

[39] R. Renger, J. A. Morin, R. Lemaitre, M. Ruer-Gruss, F. Jülicher, A. Hermann, and S. W. Grill, Co-condensation of proteins with single-and double-stranded dna, Proceedings of the National Academy of Sciences 119, e2107871119 (2022).

[40] J.-U. Sommer, H. Merlitz, and H. Schiessel, Polymer-assisted condensation: A mechanism for heterochromatin formation and epigenetic memory, Macro-molecules 55, 4841 (2022).

[41] A. Panigrahi and B. W. O’Malley, Mechanisms of enhancer action: the known and the unknown, Genome biology 22, 108 (2021).

[42] T. Quail, S. Golfier, M. Elsner, K. Ishihara, V. Murugesan, R. Renger, F. Jülicher, and J. Brugués, Force generation by protein–dna co-condensation, Nature Physics 17, 1007 (2021).

[43] M. Rubinsten, Polymer physics (United States of America, 2003).

[44] M. Gabriele, H.B. Brandão, S. Grosse-Holz, A. Jha, G. M. Dailey, C. Cattoglio, T.-H. S. Hsieh, L. Mirny, C. Zechner, and A. S. Hansen, Dynamics of ctcf-and cohesin-mediated chromatin looping revealed by live-cell imaging, Science 376, 496 (2022).

[45] E. Lieberman-Aiden, N. L. Van Berkum, L. Williams, M. Imakaev, T. Ragoczy, A. Telling, I. Amit, B. R. Lajoie, P. J. Sabo, M. O. Dorschner, et al., Comprehensive mapping of long-range interactions reveals folding principles of the human genome, science 326, 289 (2009).

[46] G. Shi and D. Thirumalai, Conformational heterogeneity in human interphase chromosome organization reconciles the fish and hi-c paradox, Nature communications 10, 3894 (2019).

[47] G. Shi and D. Thirumalai, A maximum-entropy model to predict 3d structural ensembles of chromatin from pairwise distances with applications to interphase chromosomes and structural variants, Nature Communications 14, 1150 (2023).

[48] G. Shi and D. Thirumalai, From hi-c contact map to three-dimensional organization of interphase human chromosomes, Physical Review X 11, 011051 (2021).

[49] S. Shin, G. Shi, and D. Thirumalai, From effective interactions extracted using hi-c data to chromosome structures in conventional and inverted nuclei, PRX Life 1, 013010 (2023).

