## Supplementary Information for "Active Loop Extrusion guides DNA-Protein Condensation"

### I. LANGEVIN DYANMICS SIMULATIONS

Our simulation is comprised of three types of particle: 1650 DNA particles, 5000 protein particles, and 20 SMC particles. The equation of motion of the particles is given by Langevin equation,

$$m\frac{d\mathbf{v}_i}{dt} = -\nabla U - m\gamma\mathbf{v}_i + \xi. \tag{S1}$$

Here $m$ is the mass of the particles, $U$ is the potential energy, $\gamma$ is the friction coefficient (inverse time unit), $\mathbf{v}_i$ is the velocity of $i_{th}$ particle, and $\xi$ is the delta correlated noise satisfying the fluctuation dissipation relation.

The potential energy $U$ is given by the contributions: $U = U_{\mathrm{LJ}} + U_{\mathrm{FENE}} + U_{\mathrm{ANG}}$, where $U_{\mathrm{LJ}}$ (Lennard-Jones potential), $U_{\mathrm{FENE}}$ (Finitely Extensible Nonlinear Elastic potential), and $U_{\mathrm{ANG}}$ (angular potential). We employ the Lennard-Jones potential to simulate attraction and soft repulsion interactions between DNA-Protein, Protein-Protein, and DNA-SMC. The potential is defined as:

$$U_{\mathrm{LJ}}(r_{i,j}) = 4\epsilon_{\alpha_i,\alpha_j}\Big(\big(\frac{\sigma}{r_{i,j}}\big)^{12} - \Delta_{\alpha_i,\alpha_j}\big(\frac{\sigma}{r_{i,j}}\big)^{6}\Big)\Theta(r_c - r_{i,j}). \tag{S2}$$

In this expression, $\epsilon$ and $\sigma$ denote the strength and range of the interaction, respectively. The indices $\alpha_i$ and $\alpha_j$ specify the types of particles (DNA, Protein, or SMC) for particles $i$ and $j$. The function $\Theta$ is the Heaviside step function, introducing a cut-off at $r_c = 5\sigma$. The strength and combination among different particle types in the Lennard-Jone potential are determined by the matrix $\epsilon$ and $\Delta$, respectively.

$$(\epsilon_{\alpha_i\alpha_j}) = \begin{pmatrix} \epsilon_{\mathrm{DNA,DNA}} & \epsilon_{\mathrm{DNA,PRO}} & \epsilon_{\mathrm{DNA,SMC}} \\ \epsilon_{\mathrm{DNA,PRO}} & \epsilon_{\mathrm{PRO,PRO}} & \epsilon_{\mathrm{PRO,SMC}} \\ \epsilon_{\mathrm{DNA,SMC}} & \epsilon_{\mathrm{PRO,SMC}} & \epsilon_{\mathrm{SMC,SMC}} \end{pmatrix} \tag{S3}$$

$$(\Delta_{\alpha_i\alpha_j}) = \begin{pmatrix} 0 & 1 & 1 \\ 1 & 1 & 0 \\ 1 & 0 & 0 \end{pmatrix} \tag{S4}$$

To maintain the connectivity of DNA particles, we utilize the Finitely Extensible Nonlinear Elastic potential:

$$U_{\mathrm{FENE}}(r_{i,i+1}) = -\frac{1}{2}k_F R_F^2 \log\Big[1 - \frac{(r_{i,i+1} - r^0_{i,i+1})^2}{R_F^2}\Big], \tag{S5}$$

where $k_F$ is the stiffness of the potential, $R_F$ denotes the maximum allowable displacement, and $r^0_{i,i+1}$ is the equilibrium distance between adjacent particles $i$ and $i+1$, set to $r^0_{i,i+1} = \sigma$. We set $k_F = 30\epsilon/\sigma^2$ and $R_F = 1.5\sigma$.

---

*Electronic address:
†Electronic address:

To accurately model the stiffness of DNA, we apply the angular potential:

$$U_{\mathrm{ANG}}(\theta_k) = \epsilon_b(1 + \cos\theta_k), \tag{S6}$$

where $\epsilon_b$ is the energy scale for bending. It is related to the polymer's persistence length by $\epsilon_b = k_B T l_p/\sigma$ [S1].

**Modeling loop extrusion**. To simulate the non-equilibrium, active loop extrusion (LE) process, we initiate a DNA loop upon the binding of an SMC particle to a DNA particle. This binding and subsequent loop formation are governed by the activation and deactivation of breakable harmonic potentials. The binding potential between SMC and DNA is described as:

$$U_{\mathrm{SMC,DNA}}(r_{i,j}) = -\epsilon_{\mathrm{SMC,DNA}} \log\left(1 + e^{-\frac{1}{2}k_{\mathrm{SMC,DNA}}(r_{i,j}-\sigma)^2}\right), \tag{S7}$$

where $\epsilon_{\mathrm{SMC,DNA}}$ and $k_{\mathrm{SMC,DNA}}$ are the strength and stiffness of the potential, respectively. The loop extrusion potential is given by:

$$U_{\mathrm{LP}}(r_{i,j}) = -\epsilon_{\mathrm{LP}} \log\left(1 + e^{-\frac{1}{2}k_{\mathrm{LP}}(r_{i,j}-\sigma)^2}\right), \tag{S8}$$

with $\epsilon_{\mathrm{LP}}$ and $k_{\mathrm{LP}}$ being the corresponding strength and stiffness parameters.

The loop extrusion process implemented as sequence of events, divided in three steps: 1. SMC binding, 2. Loop extrusion 3. End of the loop extrusion. Each of these steps contains three subsequent substeps as follows

1. SMC binding to DNA
   - (a) SMC binds to a DNA particle when the distance between them is less than the cutoff distance of $1.5\sigma$. Assume SMC binds at DNA index $i_{\mathrm{SMC}}$.
   - (b) Activate the binding potential $U_{\mathrm{SMC,DNA}}$ between SMC and DNA, with parameters $k_{\mathrm{SMC,DNA}} = 0.005(k_B T/\mathrm{nm}^2)$ and $\epsilon_{\mathrm{SMC,DNA}} = 100(k_B T)$. Deactivate the attractive interaction in $U_{\mathrm{LJ}}$ between SMC and DNA.
   - (c) Activate the first cross-link potential $U_{\mathrm{LP},1}$ between $i_{\mathrm{SMC}}$ and DNA particle at $i_{\mathrm{SMC}} \pm i_0$ (where $i_0 = 10$ is the initial loop size), in a randomly chosen loop extrusion direction ($\pm$). Also, activate the second potential $U_{\mathrm{LP},2}$ between $i_{\mathrm{SMC}}$ and $i_{\mathrm{SMC}} \pm i_0 \pm 1$ in the chosen extrusion direction. Initial parameters for $U_{\mathrm{LP}}$ are $\epsilon_{\mathrm{LP}} = 100(k_B T)$ and $k_{\mathrm{LP}} = 0.001(k_B T/\mathrm{nm}^2)$.
2. Loop extrusion
   - (a) Move $U_{\mathrm{LP},1}$ to the pair of beads $i_{\mathrm{SMC}} \pm 1$ and $i_{\mathrm{SMC}} \mp i_0 \mp 2$. Simulate for $n_{\mathrm{LE}}/2$ steps, where $1/n_{\mathrm{LE}}$ defines the loop extrusion speed. We set $n_{\mathrm{LE}} = 10^6$ in order for a maximum steps in the simulation to be 100 loop extrusion steps ($\sim 4\mu$m DNA loop). Parameters for $U_{\mathrm{LP}}$ remain the same.
   - (b) Move $U_{\mathrm{LP},2}$ to the pair $i_{\mathrm{SMC}} \pm 2$ and $i_{\mathrm{SMC}} \mp i_0 \mp 3$. Simulate for another $n_{\mathrm{LE}}/2$ steps. Parameters for $U_{\mathrm{LP}}$ are unchanged.
   - (c) Continue (a) and (b) until SMC unbinds from DNA, the loop breaks, or the loop extrusion reaches the determined end of DNA.
3. End of the loop extrusion
   - (a) Unbinding of SMC from DNA. If the distance between SMC and SMC bound DNA becomes greater than some threshold distance $d = 100\sigma$, then stop the loop extrusion by removing $U_{\mathrm{SMC,DNA}}$, $U_{\mathrm{LP},1}$ and $U_{\mathrm{LP},2}$.
   - (b) Breaks of the loop. If the distance between DNA particles associated with $U_{\mathrm{LP},1}$ or $U_{\mathrm{LP},2}$ becomes greater than some threshold distance $d = 100\sigma$, then stop the loop extrusion by removing $U_{\mathrm{SMC,DNA}}$, $U_{\mathrm{LP},1}$ and $U_{\mathrm{LP},2}$.
   - (c) Loop extrusion reaches to the determined end of DNA. Once the loop extrusion reaches to the end position, stop updating the position of $U_{\mathrm{LP},1}$ and $U_{\mathrm{LP},2}$.

In the Langevin dynamics simulation we use the reduced units,

$$m = \frac{k_B T \tau^2}{\sigma^2}; \quad \tau = 1/\gamma, \tag{S9}$$

where $\tau$ is the unit time in our simulation and $\gamma = 0.01\ \mathrm{ns}^{-1}$. We performed Langevin dynamics simulations using the openMM software package [S2] with the time step $\Delta t = 0.01\tau$. Simulations were carried out in a rectangular box with dimensions $2\mu\mathrm{m} \times 2\mu\mathrm{m} \times 15\mu\mathrm{m}$, implementing periodic boundary conditions.

| Parameter | Meaning | Value |
|---|---|---|
| $\epsilon_{\text{DNA,DNA}}$ | DNA-DNA interaction for LJ potential | 1 $k_BT$ |
| $\epsilon_{\text{SMC,SMC}}$ | LEF-LEF interaction for LJ potential | 1 $k_BT$ |
| $\epsilon_{\text{SMC,DNA}}$ | LEF-DNA interaction for LJ potential | 10 $k_BT$ |
| $\epsilon_{\text{PRO,PRO}}$ | Protein-Protein interaction for LJ potential | 2 $k_BT$ |
| $\epsilon_{\text{DNA,PRO}}$ | DNA-Protein interaction for LJ potential | Same as $\epsilon_{\text{PRO,PRO}}$ |
| $\sigma$ | Size of a particle | 10 nm |
| $\epsilon_b$ | Energy scale for angle potential | 5 $k_BT$ |

TABLE I: List of parameters and the values of the potential chosen in the molecular dynamics simulation.

### II. CONDENSATE GROWTH AND OSTWALT RIPENING UNDER DNA TENSION

We here discuss the dynamics of condensate growth under the tension on DNA, which leads to the enhanced Ostwalt ripening. Let us consider the situation in Fig.3a in the main text where a condensate is located outside the DNA loop. We define the difference of the chemical potential ($\Delta\mu$) between the condensate ($\mu_d$) and the stretched DNA ($\mu_p$),

$$\Delta\mu \equiv \sigma \frac{\partial \Delta F_a(L_d, L_A, L_{\text{end}}, L_c)}{\partial L_d} = \mu_d(L_d) - \mu_p(L_d, L_A, L_{\text{end}}, L_c), \tag{S10}$$

where

$$\mu_d(L_d) \equiv \sigma \frac{\partial F_d(L_d)}{\partial L_d} = \sigma \left( -v\alpha + \gamma \frac{8\pi}{3} \left(\frac{3\alpha}{4\pi}\right)^{2/3} L_d^{-1/3} \right) \tag{S11}$$

and

$$\mu_p(L_d, L_A, L_{\text{end}}, L_c) \equiv -\sigma \frac{\partial F_p(L_d, L_{\text{end}}, L_c - L_A)}{\partial L_d} = \sigma \left( -\frac{k_BT}{4l_p} \left( \frac{L_{\text{end}}^2}{(L_{\text{end}} + L_d + L_A - L_c)^2} + \frac{2L_{\text{end}}^2}{(L_d + L_A - L_c)^2} \right) \right). \tag{S12}$$

Here $\sigma$ is the length of a DNA segment which is set to 1 for simplicity. To investigate the dynamics of the condensate growth, let us write,

$$\dot{L}_d = -\gamma \Delta\mu, \tag{S13}$$

where $\gamma$ is a motility coefficient. The equilibrium is given by the condition $\Delta\mu = 0$. Eq.S13 has two fixed points for $L_d$ corresponding to the two stages of the condensate growth. First stage is the initial nucleation of protein-DNA condensate and the second stage is the growth of the condensate by further inclusion of DNA into the condensate. The first process is approximately independent to the DNA tension while the second process is tension dependent.

To illustrate the two fixed points, we start by obtaining the positions of the two fixed points. The first fixed point corresponds the initial nucleation of the condensate, which is approximately obtained by

$$v\alpha - \gamma \frac{8\pi}{3} \left(\frac{3\alpha}{4\pi}\right)^{2/3} L_d^{-1/3} = 0. \tag{S14}$$

This fixed points is unstable. The condensates smaller than the critical size diminish while the condensates larger than the critical size grow. See Fig.S1a.

The second fixed point corresponds to the condensate growth by the inclusion of DNA, which is approximately given by

$$v\alpha - \frac{k_BT}{4l_p} \left( \frac{L_{\text{end}}^2}{(L_{\text{end}} + L_d + L_A - L_c)^2} + \frac{2L_{\text{end}}^2}{(L_d + L_A - L_c)^2} \right) = 0. \tag{S15}$$

This fixed point is stable, see Fig.S1b.

Therefore the dynamics of condensate growth, Eq.S13, has two fixed points, one is unstable and the other is stable (Fig.S1c). The effect of the DNA loop, $L_A$, has significant effect to the dynamics of the condensate growth. High values of $L_A$ lead to the loss of the fixed points rendering any formation of condensates *outside* the loop unstable (Fig.S1c, see the loss of the intersection at $\dot{L}_d/\gamma = 0$ for the high values of $L_A$). Therefore when the condensate within DNA loop and outside the DNA loop coexist, the condensate outside loop tends to diminish for high $L_A$. In turn, the condensate inside the loop, which is stable, should grow instead. Thus the tension along DNA (DNA loop by loop extrusion process in our study) accelerates the dynamics of Ostwalt ripening of condensate within the DNA loop.

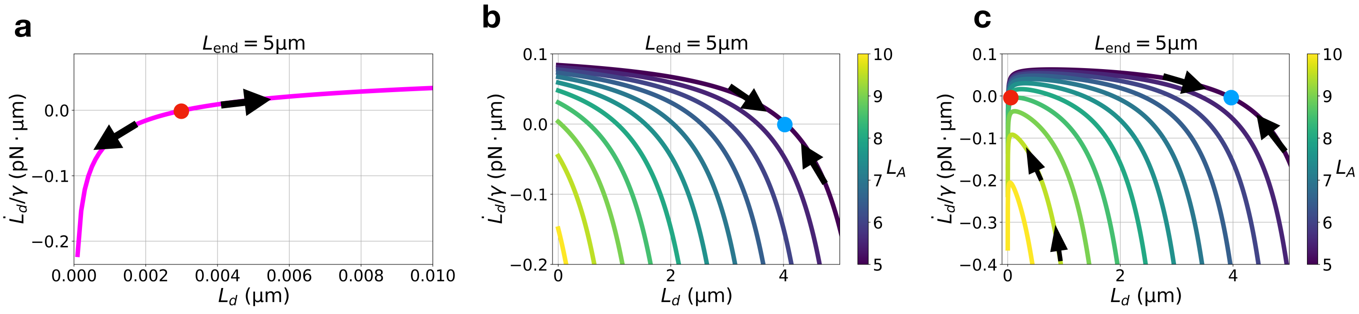


FIG. S1: (a) The first fixed point obtained from Eq.S14. The red dot indicates the unstable fixed point and the arrows are the direction of the evolution of $L_d$. (b) The second fixed point obtained from Eq.S15. The blue dot indicates an example of stable fixed point for a value of $L_A$. The arrows are the direction of the evolution of $L_d$. The color bar shows the values of the DNA loop length $L_A$ due to the loop extrusion process. (c) The dynamics of the condensate growth from Eq.S13. The red and blue dots are the unstable and stable fixed points, respectively.

### III. PROBABILITY OF CONDENSATE FORMATION

We can compute the probability of the condensation using the obtained free energies. We first computed the probability density creating DNA-protein co-condensates as a function of $L_d$. For the scenario Fig.3a in the main text,

$$P_a(L_d) = \frac{e^{-\beta\Delta F_a(L_d)}}{\int_0^{L_{\max}} dL_d e^{-\beta\Delta F_a(L_d)}}, \tag{S16}$$

where $L_{\max} = L_c - L_{\text{end}} - L_A$. The probability density has either single or double peaks ($L_d = 0$ and $L_d > 0$) depending on the parameters. The probability of the condensate formation was determined as the fraction of the peak value located at $L_d > 0$ to the sum of the peak values at $L_d = 0$ and $L_d > 0$.

### IV. ANALYSIS OF SIMULATION DATA

#### A. Condensate detection

For the identification of condensates within our simulation trajectories, we employed Density-based Spatial Clustering of Applications with Noise (DBSCAN [S3]). DBSCAN operates based on two key parameters: the distance threshold $\epsilon$ and the minimum number of points required to constitute a dense region, denoted as minPts. In our analysis, we configured these parameters to $\epsilon = 20$ nm and minPts = 30. By applying DBSCAN to the particle coordinates from the simulation trajectories, we define regions classified as condensates.

#### B. Contact Maps

To construct the contact map, we analyzed the last 1000 frames of the simulation, averaging the data across three distinct ensembles for each parameter set. A contact is defined as occurring when two particles are within 30nm of each other. The metric $I$ is calculated as the values in the contact map normalized by the total number of possible interactions ($1650^2$).


[S1] J. Midya, S. A. Egorov, K. Binder, and A. Nikoubashman, The Journal of chemical physics **151**, 034902 (2019).

[S2] P. Eastman, J. Swails, J. D. Chodera, R. T. McGibbon, Y. Zhao, K. A. Beauchamp, L.-P. Wang, A. C. Simmonett, M. P. Harrigan, C. D. Stern, et al., PLoS computational biology **13**, e1005659 (2017).

[S3] M. Ester, H.-P. Kriegel, J. Sander, X. Xu, et al., in *kdd* (1996), vol. 96, pp. 226–231.
